# Hereditary Pancreatitis Model by Blastocyst Complementation in Mouse

**DOI:** 10.1101/457978

**Authors:** Ayumu Asai, Masamitsu Konno, Jun Koseki, Koichi Kawamoto, Takahiro Arai, Taroh Satoh, Hidetoshi Eguchi, Masaki Mori, Yuichiro Doki, Hideshi Ishii

## Abstract

**Summary Statement:** The present study is the first report for reproducing human disease by utilizing blastocyst complementation method and would lead to the development of novel therapy for human disease.

The application of pluripotent stem cell is expected to contribute to the elucidation of the unknown mechanism of human diseases. However, in vitro induction of cells in several organs, such as the pancreas and liver, remains difficult; therefore, reproduction of those diseases in a model has not been feasible. To reproduce human hereditary pancreatitis (HP), which is most frequently caused by the mutations in the *PRSS1* gene, we performed the blastocyst complementation (BC) method. In the BC method, mouse embryonic stem (ES) cells harboring CRISPR/CAS9-mediated mutations in the *Prss1* were injected into the blastocysts of deficient *Pdx1* gene mice, which is a critical transcription factor in the pancreas. The results showed that the blastocysts injected into the *Prss1*-mutant ES cells induced trypsin activation. This implied that the mouse phenotype mimics that of human HP and that the BC method was useful for the reproduction and study of pancreatic disorders. The present study opens the possibility of investigating uncharacterized human diseases by utilizing the BC method.

## Introduction

Induced pluripotent stem cell (iPSC) technology enables reprogramming of multipotent undifferentiated cells that can be used for the generation of disease-specific germline mutations harboring somatic cells, as well as normal healthy cells. Therefore, disease-specific iPSCs are regarded as important tools to monitor disease processes, to explore drug screening and discovery, and to elucidate the pathophysiology of human diseases (Bellin et al., 2012; Maehr et al., 2009). Previous studies have used disease-specific iPSCs from patients with hereditary disorders to clarify the pathogenesis of muscle dystrophy (Shoji et al., 2015) and fibrodysplasia ossificans progressive (Matsumoto et al., 2013). Although the use of iPSCs was successful in the reproduction of the phenotypes of those diseases, its application for pancreatic diseases, such as pancreatitis and diabetes (Borowiak and Melton, 2009), and some other diseases of the kidney (Mae and Osafune, 2015) and liver (Sakiyama et al., 2017) remains to be successfully performed, presumably due to the difficulty of cell differentiation in vivo and the complexity of the organ structures. In order to reproduce the diseases that are derived from tissues, in which induction of differentiation is difficult, it is undoubtedly necessary to improve the efficiency of differentiation induction and to reconstruct the complexity of tissues in a model.

The blastocyst complementation (BC) method has been reported to be applicable for the construction of pancreatic tissues. Mice without the knockout allele of the pancreatic and duodenal homeobox 1 (*Pdx1*) gene, which is a critical transcription factor that determines the fate of pancreatic endocrine and exocrine differentiation, have been reported to manifest with lethal phenotypes immediately after birth due to a defective formation of the pancreas, and injection of wild-type embryonal stem (ES) cells in a knockout blastocyst of the *Pdx1* gene rescued the mice completely (Offield et al., 1996; Kobayashi et al., 2010). In a previous study, the BC method was applied for pancreatic formation in an intercross species condition (i.e., rats to mice or vice versa) (Kobayashi et al., 2010). This indicated the possible human application of this regenerative medicine technology in the future.

Although the BC method using disease-specific pluripotent stem cells had been suggested to reproduce hereditary diseases in vivo, its application on the study of hereditary diseases has been reported by only a few. In the present study, we aimed to reproduce hereditary pancreatitis (HP) using the BC method. Several causative genes of HP have been reported (Applebaum-Shapiro et al., 2001; Howes et al., 2004; Rebours et al., 2009), but one sequencing study indicated that mutation in the cationic trypsinogen or protease serine 1 (*PRSS1*) gene is one of the most common causes of HP (Whitcomb et al., 1996). In this study, we established the ESCs harboring the *Prss1* mutation and successfully performed the BC method to reproduce the HP phenotype of in mice.

## Results

### Establishment of a disease-specific iPS model using mouse embryonic stem cell

Mutation in the *PRSS1* gene is known as one of the most common causes of HP (Whitcomb et al., 1996). We examined the mutated gene, the major mutation residues, and the mutation rate in HP. As a result, *PRSS1* was the most frequently mutated gene in HP (Table S1). In general, translation of the PRSS1 leads to the production of the precursor of trypsinogen, which is cleaved off from the region of its signal peptide and the trypsinogen-activating peptide, resulting in the activation of trypsin; on the other hand, cleavage of activated trypsin causes inactivation. Substitution of amino acids due to mutations, which play a role in the activation of the trypsinogen precursor or the activated trypsin, was reported to result in pancreatitis (Gorry et al., 1997). Moreover, the substitution of alanine to valine at position 16 (A16V) was reported to result in the abnormal processing of the trypsinogen precursor; whereas mutations of the *D16A*, *D22G*, and *N29I*, or *N29T* caused abnormalities in the activation of trypsin (Raphael and Willingham, 2016). Mutations in the *R122H* or *R122C* have been known to interfere with the inactivation of activated trypsin (Raphael and Willingham, 2016). The *N29I* was observed as the most common mutated gene in HP (Gorry et al., 1997), suggesting its critical role in HP. However, the precise mechanism for disease development remains to be elucidated.

To establish the disease-specific iPSCs that correspond to HP in an animal model, we compared the *PRSS1* genes in mice and humans and found 95% similarity and 76% identical amino acid sequences (Figures 1A), indicating conservation of the peptide sequence of the PRSS1 protein among the species. On the other hand, the 29th amino acid residue of PRSS1, which causes HP, was different between human (H) and mouse (T). We created both PRSS1 structures and investigated for changes after mutation of the 29th amino acid residue. As a result, each structure was destabilized by the mutation in the 29th amino acid residue (Figure 1B). Moreover, the phenotype of HP might depend on the original residue if the human PRSS1 mutation is mimicked in a mouse model. Therefore, we created two three-dimensional (3D) structures by exchanging the 29th residue in the PRSS1 of human and mouse and comparing these with the structure of each wild PRSS1. As a result, none of the residues affected the original protein structure, suggesting that changing the structure by mutating to isoleucine did not depend on the original residue (Figure 1C). We then designed the guide RNA for the introduction of T29I mutation by CRISPR/CAS9 technology (Figure 2A). By introduction of the guide RNA into mouse ESCs, we established PRSS1-mutant ESCs, the sequence of which verified introduction of the mutation in the ESCs (PRSS1^T29I^ ESCs; Figure 2B). The PRSS1^T29I^ ESCs and wild-type ESCs were cultured and maintained successfully in an undifferentiated state in a medium that contained leukemia inhibitory factor (LIF) (Figure 2C).

**Figure 1.**
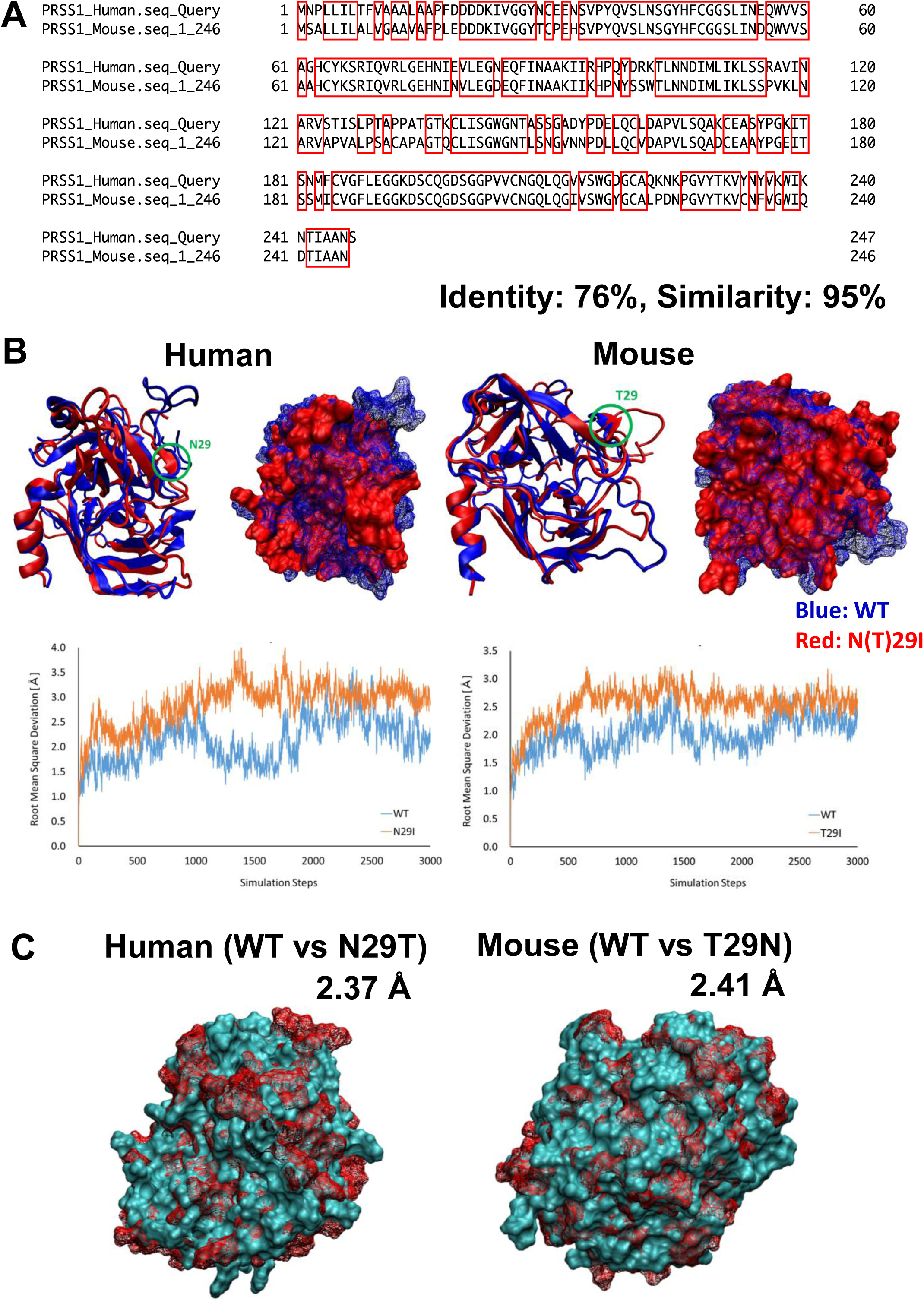
Comparison of PRSS1 between human and mouse. (A) Comparison of sequences of PRSS1 between human and mouse. Regions surrounded by red line are common residues between human and mouse. (B) Comparison of 3D structure of Prss1 between WT and N29T (human) or T29N (mouse). The difference between the two structures is represented by root mean square deviation (RMSD). (C) Comparison of 3D structure of Prss1 between WT and N(T)29I in human and mouse. Comparison of flexibility between WT and N(T)29I in the most flexible region (human: residues 146-156, mouse: residues 187-200) by thermal vibration. Vertical axis: RMSD, Horizontal axis: simulation steps.

**Figure 2.**
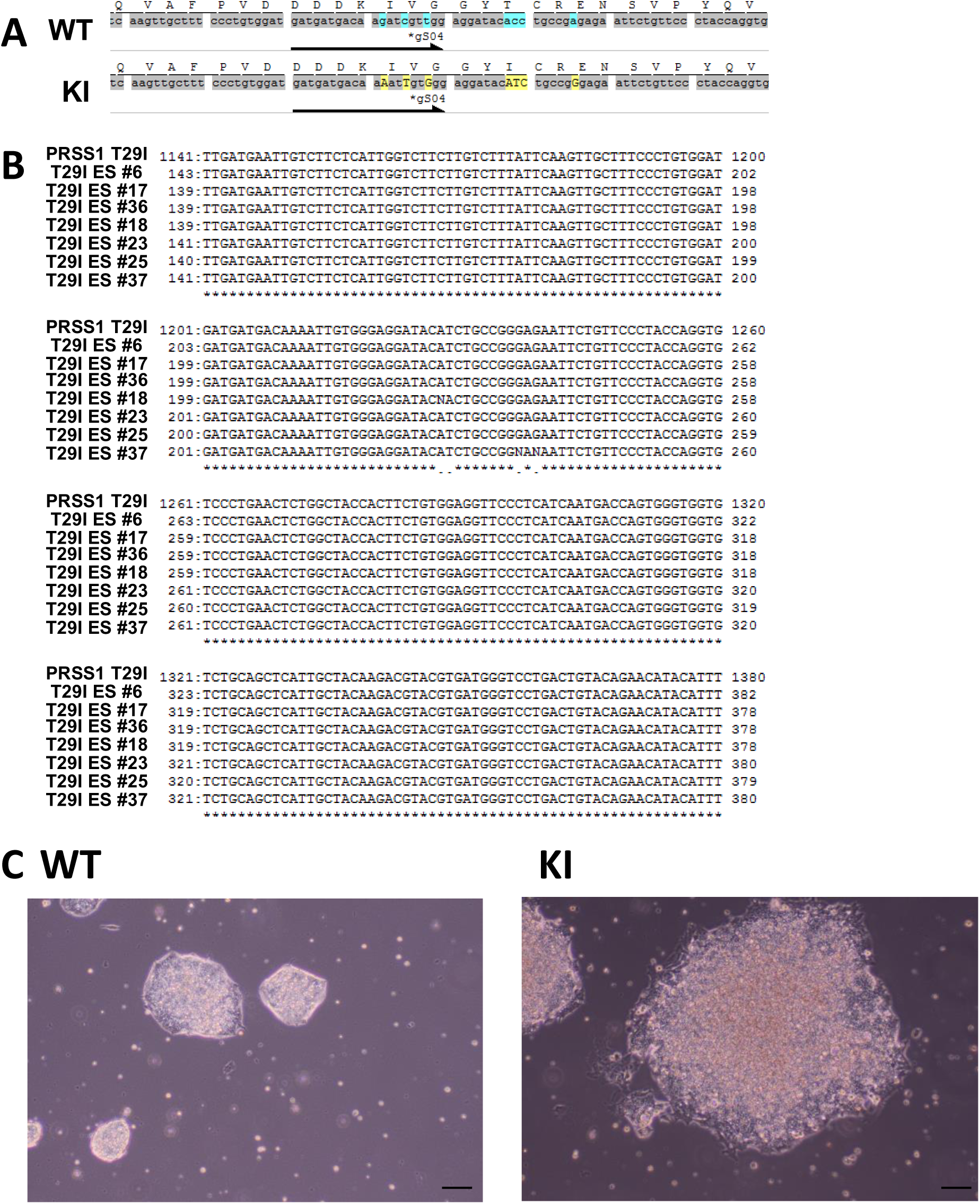
Generation of PRSS1^T29I^ ESC. (A) The sequence of *PRSS1* in wild-type mice ESCs and PRSS1^T29I^ ESCs. (B) The sequence of mouse ES cells after the introduction of mutation. (C) Phase contrast microscope images of ESCs. The WT on the left side and the PRSS1^T29I^ on the right ES indicate the colony formation of undifferentiated states in the medium with LIF. Scale bar, 200 μm.

PRSS1^T29I^ ESCs were established by CRISPR/CAS9 technology but needed to have chimera-forming ability and wild-type ESCs to reproduce HP by the BC method. First, the expression of PRSS1 in each tissue was examined using BioGPS to investigate the influence of PRSS1 mutation on the whole body. As a result, *PRSS1* was expressed in the pancreas alone in both human and mouse (Figure S1). Therefore, the PRSS1 mutation would affect only the pancreas. Second, the gene expression profiles were compared between ESCs to evaluate the chimera-forming ability of PRSS1^T29I^ ESCs. As a result, the expression of each differentiation marker was similar between PRSS1^T29I^ ESCs and wild-type ESCs (Figure 3A). Furthermore, the gene set enrichment analysis (GSEA) showed that the gene sets involved in both differentiation and stem cells were not enriched (Figures 3B and 3C). The most enriched gene set between PRSS1^T29I^ and wild-type ESCs was the set involved in DNA polymerase (Figure S2). The use of CRISPR/CAS9 technology might have affected DNA synthesis, but an enrichment score of 0.57 was not much. Therefore, the PRSS1^T29I^ would retain its chimera-forming ability.

**Figure 3.**
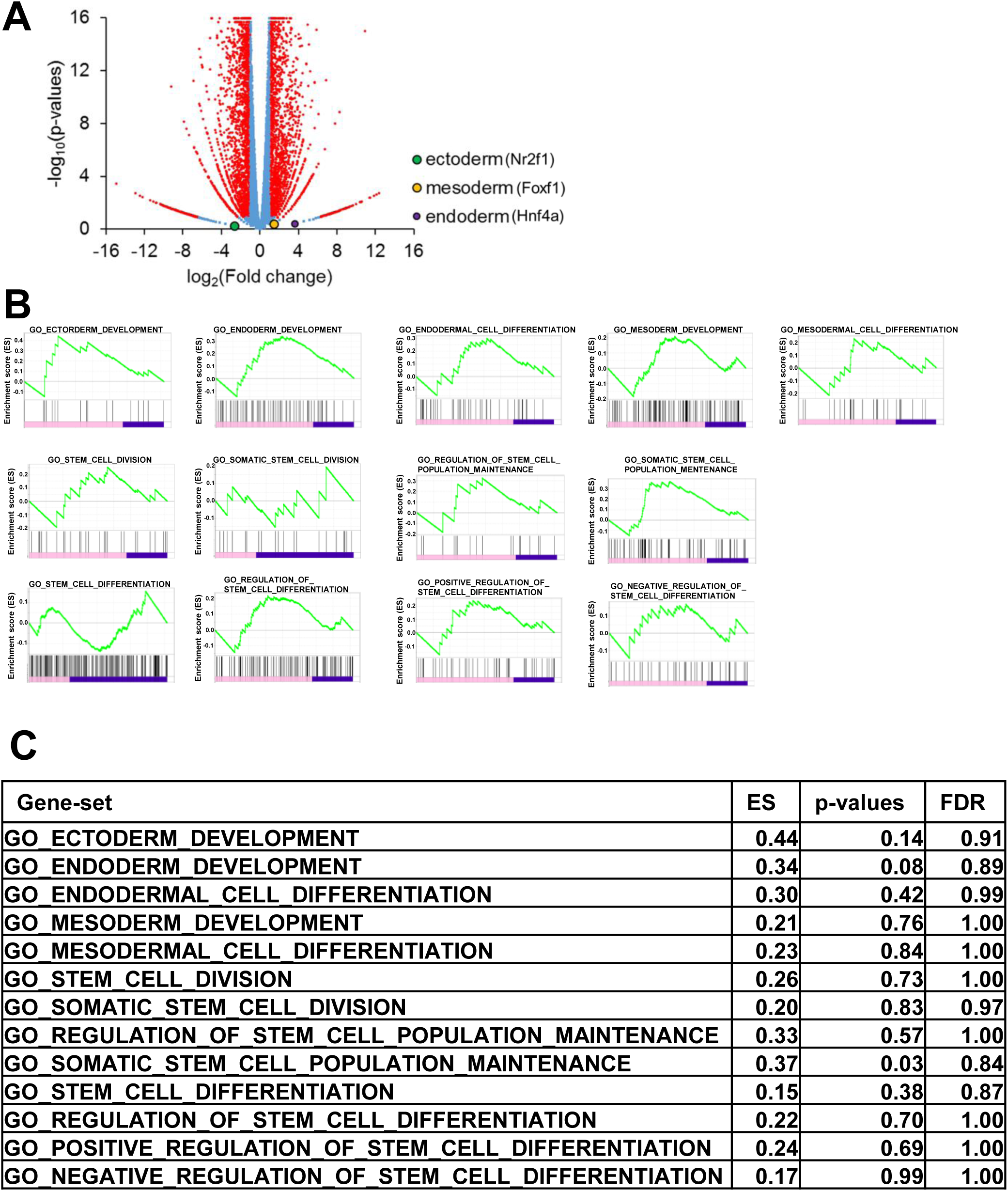
Gene expression of PRSS1^T29I^ ESCs. (A) Volcano plot for the gene expression of PRSS1^T29I^ against wild-type ESCs. Horizontal axis: logarithm of fold change in the gene expression for PRSS1^T29I^ against wild-type ESCs. Vertical axis: logarithm of p-values. (B) Enrichment plot of the gene sets involved in differentiation and undifferentiation. (C) ES, p-values, and false discovery rate of the gene sets involved in differentiation and undifferentiation.

### Reproduction of HP using disease-specific pluripotent stem cells

The *Pdx1*-null mice that lacked pancreas were dead at two or three days after birth. However, introduction of wild-type ESCs in *Pdx1*-deficient ESCs was reported to result in the complete rescue of the lethal phenotypes that harbored pancreatic tissues, with 100% contribution of wild-type ESCs. In the field of regenerative medicine, the BC methods have emerged recently organ formation. Here we applied the BC method for the contribution of disease-specific PRSS1^T29I^ ESCs and the causative formation of pancreatitis. In this experiment, we injected PRSS1^T29I^ ESCs into the blastocysts that were derived from *Pdx1*-null mice and were transferred to another female, in which pregnancy was mimicked by injection of human chorionic gonadotropin (hCG), creating a provisional belly. Although a newborn was successfully delivered, it died 1.5 days after birth (Figure 4A).

**Figure 4.**
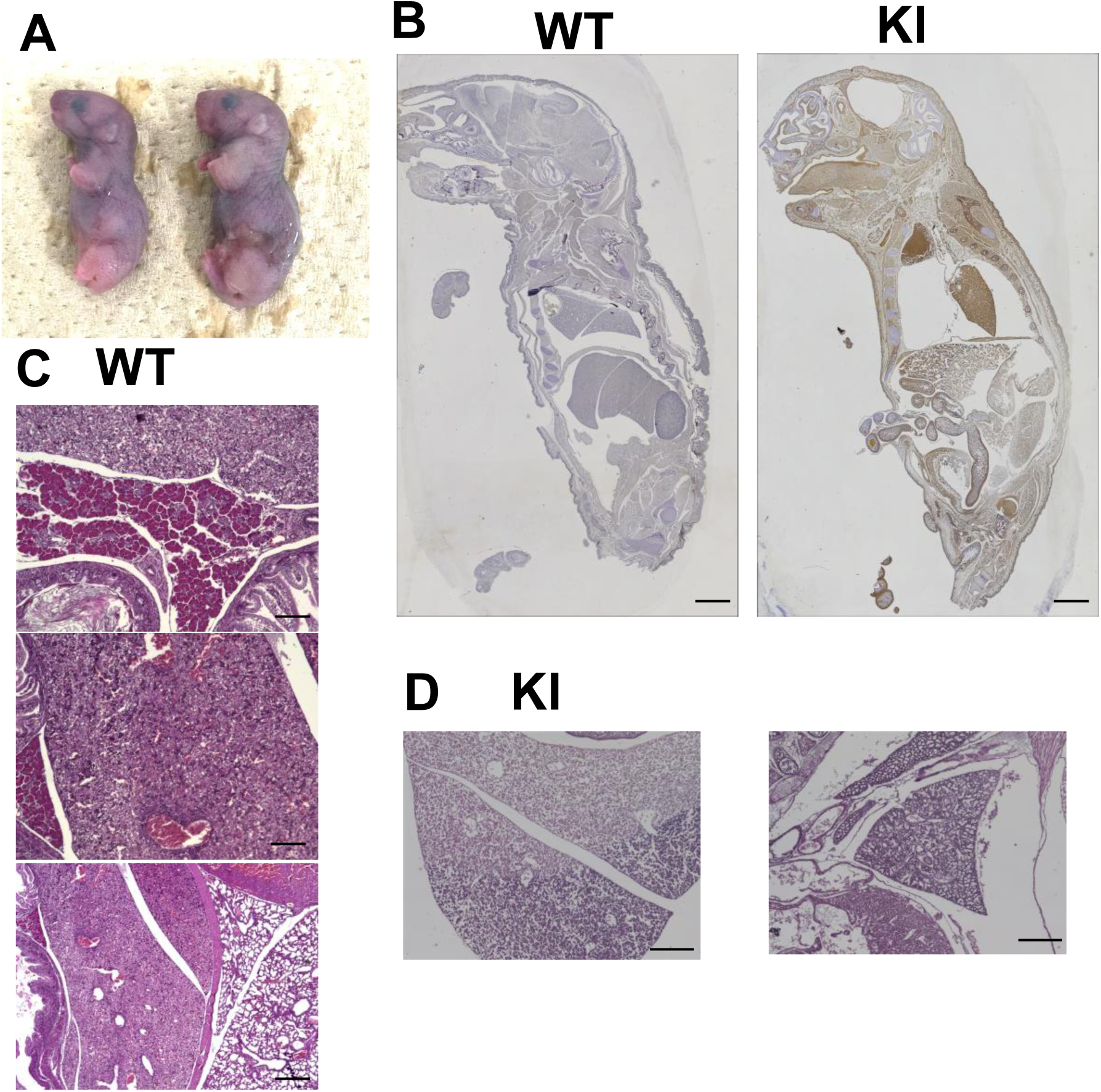
Reproduction of HP. (A) Image of newborn *Pdx1*-KO mice on day 1 after being injected with PRSS1^T29I^ ESCs. (B) The PRSS1^T29I^ ESCs are Gfp-positive, which is shown as a brown chromogen color. Scale bar, 200 μm. (C) The hematoxylin and eosin (HE) staining of wild-type mice at E19.5. Scale bar, 20 μm. (D) The HE staining of *Pdx1*-KO mice that were injected with PRSS1^T29I^ ESCs at E19.5. Scale bar, 50 μm.

To study the situation of PRSS1^T29I^ ESCs-derived from the embryo in the uterus, we decided to sacrifice the pregnant mice on E19.5 days. We were interested to know whether the injected PRSS1^T29I^ ESCs contributed to the tissues of the pancreas. We performed an anti-Gfp antibody immunostaining experiment on E19.5-day embryos, because the PRSS1^T29I^ ESCs were labeled by the *GFP* gene. The results indicated that Gfp-positive cells were detected in the entire pancreas and in the extra-pancreatic organs, such as the liver (Figure 4B). These suggested that the injected PRSS1^T29I^ ESCs contributed to the formation of chimera. Histologic examination of the embryos derived from the injection of PRSS1^T29I^ ESCs showed no apparent formation of pancreatic organs (Figure 4D), whereas those from wild-type ESCs had formed a pancreas (Figure 4C). Moreover, the liver of PRSS1^T29I^-derived embryos showed severely destroyed structures and thin diaphragm (Figure 5A), presumably due to the effects of activated trypsin. We then measured the trypsin activity in mice tissues of PRSS1^T29I^ ESCs-derived and wild-type ESCs-derived embryos. The results indicated that the trypsin activity in PRSS1^T29I^ ESCs-derived embryos was significantly higher than that in wild-type ESCs-derived embryos (Figure 5B). The present study showed that the BC methods of PRSS1^T29I^ ESCs can reproduce the phenotype of HP in a model.

**Figure 5.**
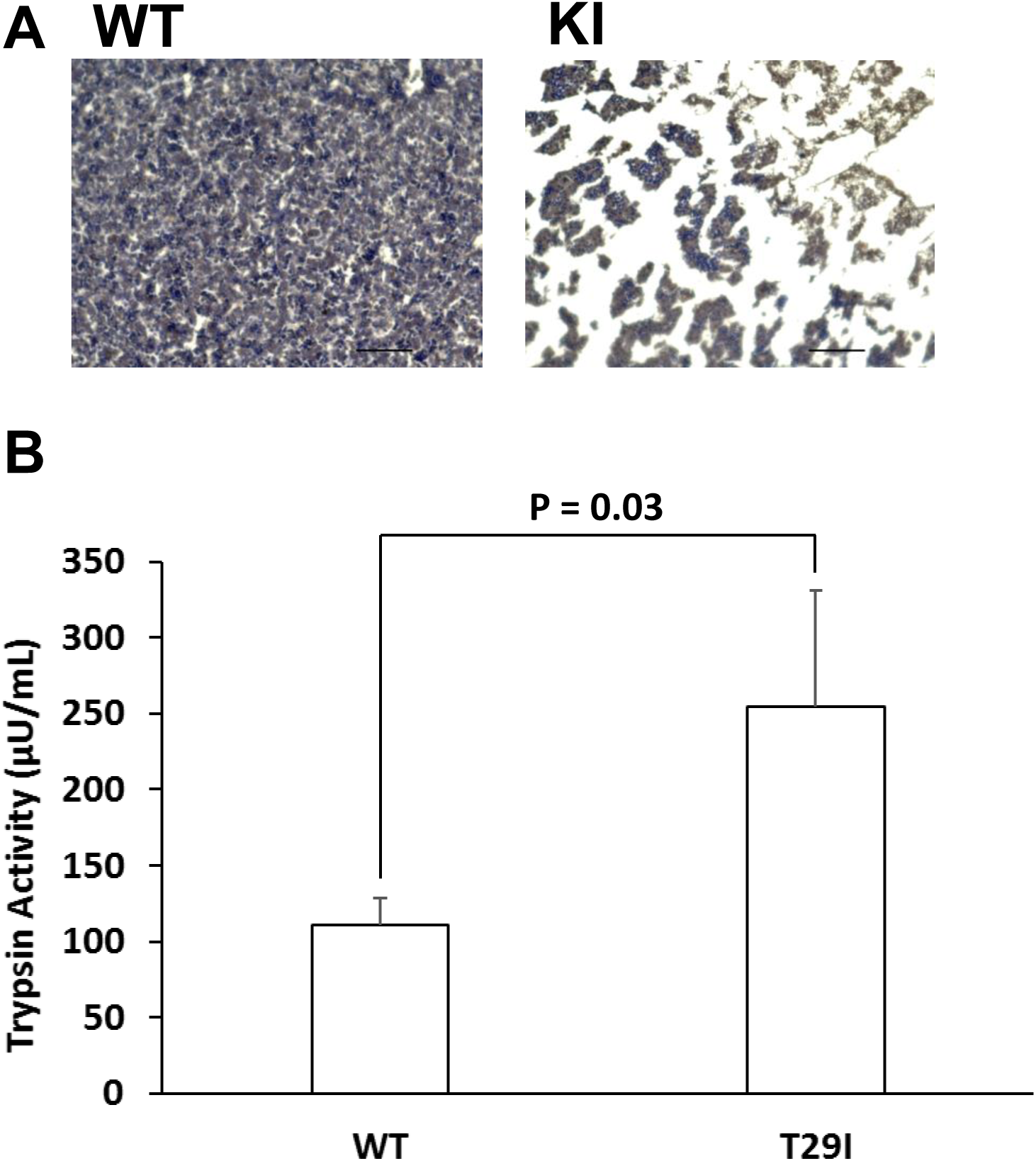
Trypsin activity of the HP model. (A) The HE staining of the liver of the wild-type mice and the *Pdx1*-KO mice that were injected with PRSS1^T29I^ ESCs at E19.5. Scale bar, 30 μm. (B) The results of the measurement of trypsin activity. Each value represents the mean ± S.D. (n = 3).

## DISCUSSION

HP is a relatively rare disorder of the pancreas (Applebaum-Shapiro et al., 2001; Howes et al., 2004; Rebours et al., 2009) and was first reported by Comfort and Steinberg in 1952 (Comfort and Steinberg, 1952). HP is associated with pancreatic inflammation and had been attributed to genetic causes, such as mutation in the long arm of chromosome 7q35 of the *PRSS1* gene, which was the first isolated gene responsible for HP (Whitcomb et al., 1996). The symptom and etiology of HP follow a similar pattern with those of alcohol-associated chronic pancreatitis, but HP shows a distinct phenotype of earlier onset of pancreatitis in a younger age, compared with the later stage onset in the disease progression of diabetes mellitus (Howes et al., 2004). Recurrent pancreatitis with severe abdominal pain could be refractory to conventional non-steroidal anti-inflammatory drugs and could interfere with social activities; moreover, severe cases are subjected to surgical resection of the inflamed regions (Howes et al., 2004). Inflammation with infiltration of lymphocytes and neutrophils can damage cells and lead to malignant transformation, and one European study indicated an increasingly high risk of pancreatic cancer unrelated to the genotype after the age of 50 years (Howes et al., 2004). Therefore, further study on HP-specific iPS cells that were harvested from patients’ cells would be needed to elucidate the mechanism and for drug screening. Moreover, the correlation of pancreatic duct inflammation with epithelial destruction, regeneration, and cellular transformation remains to be understood. Given that *PRSS1* gene mutation is a major cause of HP, which is refractory and is frequently associated with tumors, designing effective therapeutic strategies through novel cell culture technologies, animal models mimicking human disease, and pain management tools is necessary (Uc et al., 2016).

Because *PRSS1* was the most frequently mutated gene in HP, we established the corresponding animal model. The sequence study indicated that asparagine was substituted to isoleucine at position 29 (N29I) in human, although a previous study indicated that a transgenic mouse that overexpressed the amino acid substitution of threonine to isoleucine at position 29 (T29I) showed the phenotype of recurrent acute and chronic pancreatitis (Gorry et al., 1997), through the involvement of apoptosis and necrosis of pancreatic cells (Kaiser et al., 1995; Bhatia et al., 1998; Mareninova et al., 2006). Furthermore, overexpression of human N29I in mouse cells resulted in the induction of apoptosis of murine acinar cells, which is of a similar phenotype with that of pancreatitis (Athwal et al., 2014). These data indicated that N29I in human and T29I in mouse have similar effects on the onset of pancreatitis. Similarly, the present study indicated that the computational predicted structure of T29I was similar with that of N29I. In this study, we used the CRISPR/CAS9-mediated mutation of T29I substitution in the endogenous copy of the *Prss1* gene allele in a mouse BC model, suggesting that T29I was a proper counterpart of human PRSS1^N29I^ mutation. However, we have to consider the possibility that mouse PRSS1^T29I^ mutation might express a more severe phenotype than human PRSS1^N29I^, based on the fact that the mice died immediately after birth.

A recent report indicated that a 3D human iPS-cell engineered heart tissue was a useful tool for modeling *torsade de pointes*, which is a lethal arrhythmia that is often drug-induced; in particular, the 3D model provided details on the mechanisms underlying arrhythmia generation and on the means for drug discovery and safety tests (Kawatou et al., 2017). Moreover, some studies have reported the usefulness of patient-derived iPSCs as screening biomarkers for Alzheimer’s disease (Shirotani et al., 2017) and Behçet’s disease (Son et al., 2017). The use of iPSCs and animal model is expected to cause a breakthrough in the discovery of drugs that target the development of rare genetic disorders, which accounts for approximately 7,000 diseases that affect millions of individuals in the United States (Sun et al., 2017). Given that disease-specific iPS cells are in the undifferentiated state, in which alterations in the disease-causing gene sets are preserved (Rebours et al., 2009), a technology that induces differentiation of the disease-specific iPS cells would be expect to reproduce the initial process of disease before its onset.

A recent study indicated that the BC method was applicable to intercross species of mice ES-in-rats blastocysts as well as rats ES-in-mice blastocysts (Kobayashi et al., 2010); also, it has been demonstrated that swine ES-in-swine blastocysts (Matsunari et al., 2013), as well as mice ES-in-rats blastocysts were useful to maintain the insulin level in glucose metabolism, as experimental models (Yamaguchi et al., 2017). Furthermore, knockout of endoderm-specific gene *Mixl1* indicated that targeted organ generation using *Mixl1*-inducible mouse pluripotent stem cells in BC was able to induce somatic organs, such as the gut, liver, and pancreas (Kobayashi et al., 2015).

In the present study, we established PRSS1^T29I^ ESCs. The successful maintenance of the culture of PRSS1^T29I^ and wild-type ESCs in a medium containing LIF suggested that the PRSS1^T29I^ ESCs were in the undifferentiated state. Moreover, the gene expression profiles involved in the differentiation and undifferentiation of each ESC were confirmed by RNA-seq to be similar between the ESCs, suggesting that the chimera-formation ability was similar between PRSS1^T29I^ ESCs and wild-type ESCs. On the other hand, GSEA revealed that the gene set involved in DNA polymerase activity was the most enriched. Although introduction of a mutant gene by CRISPR/CAS9 might have affected DNA polymerase, the enrichment score was not so high and the PRSS1^T29I^ ESCs were maintained in the undifferentiated state. Therefore, we considered that the result of PRSS1^T29I^ of did not affect the undifferentiated state and chimera-formation ability of PRSS1^T29I^ ESCs. In other words, establishment of ESCs by *Prss1* mutation succeeded without affecting the chimera-formation ability.

We applied the BC method for PRSS1^T29I^ ESCs to reproduce human HP. In the mice in which the BC method was applied, the PRSS1^T29I^ ESCs contributed to the formation of the whole pancreas and the chimera. Subsequently, the mice-derived PRSS1^T29I^ ESCs resulted in activation of trypsin and destroyed liver and lung. These results indicated that the BC method for the PRSS1^T29I^ ESCs succeeded to reproduce the pathology of human HP (Figure 6). However, the mice died immediately after birth and more severe disease onset than that in human. Further improvement of the BC method is necessary to elucidate disease pathophysiology and drug screening. In present study, BC method was able to reproduce pancreatic disease and is expected to reproduce disease in other organs.

**Figure 6.**
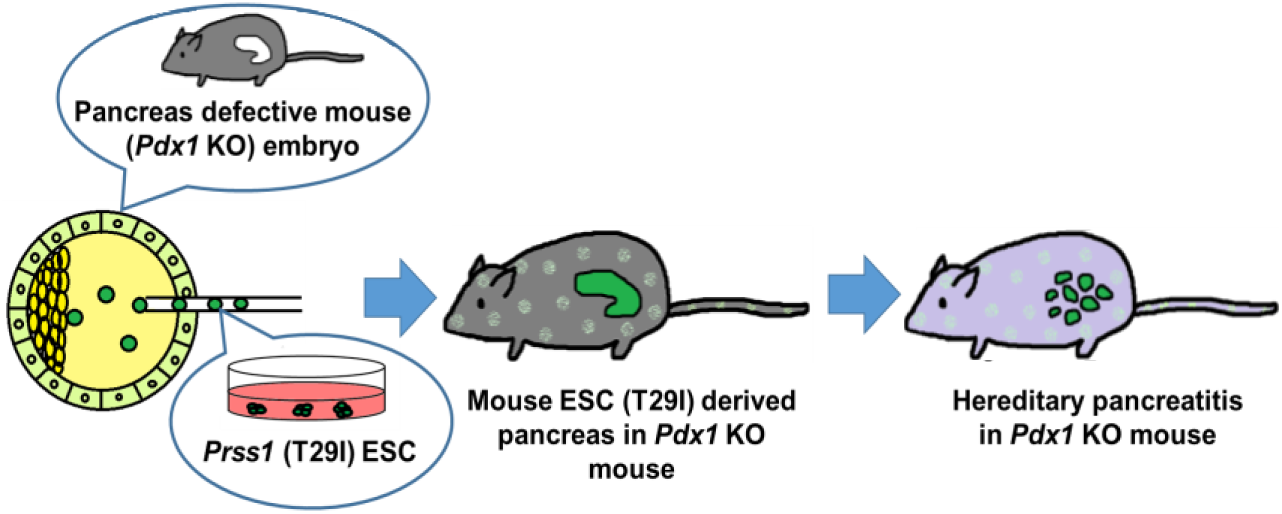
Reproduction of HP by BC method. Mouse ES cells harboring mutations in the *Prss1* were injected into the blastocysts with deficient *Pdx1* gene. The blastocysts injected into the *Prss1*-mutant ES cells induced trypsin activation like human HP.

In the present study, we were able to reproduce HP by the BC method. The embryos derived from PRSS1^T29I^ ESCs, not from wild-type ESCs, showed features of extensive digestion by trypsin in the pancreatic lesion, as well as in the liver and lung of the *Pdx1*-deficient blastocyst body. This suggested the contribution of the transcription factor *Pdx1* in a broad range of organs, which is compatible with the intercross species BC method (Kobayashi et al., 2010). Although the application of this method to human cells has ethical issues to be overcome, the use of disease-specific iPS cells would be an attractive tool for the elucidation of disease mechanism and can contribute to the progression of medicine.

## Materials and Methods

### Protein structure analysis

We created the 3D structures of wild-type (WT) PRSS1 and mutated PRSS1 (human: N29I, mouse: T29I) using the homology modeling method, based on the crystal structure of the human cationic trypsin G193R mutant (PDB ID: 4WWY). Then, the structures of the WT and mutated type were subjected to energy minimization in the water phase of the AMBER force field using the AMBER12 program package. Using these minimized structures, we performed molecular dynamics (MD) simulation to investigate the differences in flexibility between the WT and mutated structures. We performed the MD simulation (i.e., elevated temperature process and thermodynamically conformational sampling) at around 37 °C (310 K) using the periodic boundary condition. The flexibility of each protein structure was compared by calculating the root mean square deviation (RMSD) from each initial coordinate. Moreover, we created the structures that were exchanged at the 29th amino acid residue in PRSS1 between humans (N→T) and mice (T→N) and compared the original structures with the mutated structures. The RMSD was calculated as an index of structural difference in each minimized structure after *in silico* annealing processes.

### Establishment of a disease-specific iPS model using mouse embryonic stem cell

PRSS1^T29I^ ESCs were generated using using gRNAs in the CRISPR/CAS9 System (Figure 1A). On the first day, pregnant male serum gonadotropin (5 IU) was administered intraperitoneally to female *Pdx1* KO hetero C57BL/6J mice. On the third day, human chorionic gonadotropin (5 IU) was administered intraperitoneally to the mice. Subsequently, the mice were mated with male *Pdx1* KO hetero C57BL/6J mice. On the sixth day, blastocysts were collected from the female mice and were injected with PRSS1^T29I^ ESCs. Subsequently, the blastocysts were cultured overnight. On the seventh day, the cultured blastocysts were transplanted into the uteri of the foster mothers.

### ESC culture

The v6.5 mouse ES cells, which were derived from a male mouse embryo, were cultured in Dulbecco’s modified eagle medium (Nakalai Tesque, Japan) containing 15% FBS (Thermofisher); 1% non-essential amino acids (NakalaiTesque, Japan); 1% sodium pyruvate (Nakalai Tesque, Japan); and LIF (NakalaiTesque, Japan) at 37 °C in a humidified atmosphere with 5% CO_2_.

### RNA-seq for ESCs

RNA was extracted from each ESC and was sequenced by Hi-seq2500. Gene expression was analyzed using CLC Genomics Workbench Ver.9.0 and GSEA software.

### Immunostaining

Immunohistochemical analysis was performed on 3.5-μm, paraffin-embedded sections from the xenografts. The paraffin-embedded sections were de-paraffinized in Hemo-De (Farma, Japan) and were rehydrated in a graded series of ethanol. The slides were heated in an antigen retrieval buffer for 40 minutes, blocked with goat or horse serum for 20 minutes at room temperature, and incubated with monoclonal mouse anti-GFP antigen antibody (1:1000, Abcam, England) overnight at 4 °C. The Vectastain ABC System (Vectastain, Funakoshi, Japan) was used to visualize the antigens. Counter-staining was performed using hematoxylin only or HE.

### Trypsin activity assay

Trypsin activity was evaluated using a Trypsin Activity Assay Kit (ab102531, Abcam). The abdomen of each mouse at birth (day 0) was cut and was measured as a sample.

## Acknowledgments

We thank the members of our laboratories.

## Data availability

The sequence data have been deposited in the Gene Expression Omnibus under ID codes GSE119172.

